# Mechanical collaboration between the embryonic brain and the surrounding scalp tissues

**DOI:** 10.1101/2021.05.05.442865

**Authors:** Koichiro Tsujikawa, Kanako Saito, Arata Nagasaka, Takaki Miyata

**Author notes:** These authors contributed equally to this work (Koichiro Tsujikawa, Kanako Saito). Corresponding author: Takaki Miyata.

## Abstract

During brain enlargement between embryonic day (E) 11 and E13, within the limited mouse cranial space enclosed by the scalp consisting of epidermal and preosteogenic mesenchymal layers, the angle between the pons and the medulla decreases. This raises the possibility that the scalp, upon being pushed outwardly by the brain and stretched, in return inwardly recoils to confine and fold the brain. Our stress-releasing tests showed that the scalp recoiled to shrink more extensively at E12~13 than at E15~16 and that the *in vivo* pre-stretch prerequisite for this recoil response depended physically on the brain (pressurization at 77~93 Pa) and on actomyosin and elastin within the scalp layers. Under non-stretched conditions, scalp cell proliferation declined, while re-stretching of the shrunken scalp rescued proliferation. In scalp-removed heads, pons–medulla folding was reduced, and the spreading of ink from the lateral ventricle to the spinal cord that occurred in scalp-intact embryos (with >5 μl injection) was lost, suggesting that the scalp plays roles in brain morphogenesis and cerebrospinal fluid homeostasis. Thus, the brain and the scalp mechanically interact and collaborate.

## INTRODUCTION

In a variety of morphogenetic events, inter-tissue forces play important roles (Goodwin and Nelson, 2021). For example, mechanical influences on blastocysts from the trophectoderm (Weberling and Zernicka-Goetz, 2021) or those on cylinder-shaped embryos from the uterine wall (Hiramatsu et al., 2013) facilitate embryogenesis. Additionally, the development of tubular structures, such as the gut (Savin et al., 2011; Shyer et al., 2013), bronchi (Kim et al., 2015; Varner et al., 2015), and oviducts (Koyama et al., 2016), is driven by mechanical relationships between the inner and outer layers, enabling villification, branching or fold pattern formation of the inner epithelial sheet. In these examples, constraint or confinement by an external or superficial layer is a key mechanical factor that enables the deeper structure to continue normal development (Nelson, 2016; Trushko et al., 2020). In the developing mammalian brain, intramural cellular and tissue mechanics have recently been explored (Okamoto et al., 2013; Koser et al., 2016). However, how embryonic brain vesicles mechanically interact with the outer surrounding mesenchymal and epidermal tissues and the biological significance of these interactions remain unknown. Through “morphomechanical” approaches combining experimental and computational data, Taber and colleagues found that invagination during chicken optic cup formation is driven by external confining factors such as the ectoderm and the extracellular matrix (Hosseini et al., 2014; Oltean et al., 2016). A sagittal sectional survey of how brain vesicles grow under space limitation within the heads of mouse embryos (Fig. S1A, B) showed that the angle between the pons and the medulla dramatically decreases between embryonic day 11 (E11) and E13, thereby narrowing the fourth ventricle. We accordingly hypothesized that the dorsal “scalp” (herein designated on the basis of gross anatomy but histologically consisting of an epidermal layer and the underlying mesenchyme) might play a role in confinement of the growing brain, inducing brain folding to decrease the pons–medulla angle (Fig. S1B). To investigate this possibility, we firstly investigated the histological and mechanical properties of the scalp, and examined brains’ responses upon removal of the scalp. The mesenchymal element of the calvarial (i.e., dorsally brain-covering) connective tissues during this E11–E13 period does not yet stain with Alizarin red or alkaline phosphatase and is known to be preosteogenic (Morriss-Kay, 2001; Rice et al., 2003; Roybal et al., 2010; Cesario et al., 2018). Since this mesenchymal layer and the adjacent overlying epidermal layer both tangentially expand as the brain grows outwardly, we also investigated whether possible mechanical influences from the expansively growing brain promote the development of the scalp (Fig. S1B).

## RESULTS

### Immunohistochemical characterization of the calvarial epidermal and mesenchymal tissues during the E11–E13 period

We first subjected coronal sections of the heads of E12 embryos to immunohistochemistry using antibodies as listed in Fig. 1. Low-magnification observation showed that superficial immunoreactivity for alpha smooth muscle actin (αSMA), which has been suggested to function in cells playing force-generating and/or constricting roles (Shyer et al., 2013; Kim et al., 2015; Miyai et al., 2019; Haddem et al., 2020), was continuously strong throughout the dorsal (calvarial) side of the head but not ventrally towards the face, producing a staining pattern similar to a cap (Fig. 1A). High-power views of the dorsal scalp (Fig. 1B) revealed that the most superficial layer was positive for cytokeratin 10 (CK10); therefore, this layer was identified as epidermis. In a deeper zone, fibre-like immunoreactivity for αSMA was abundantly found, and most cell nuclei were positive for Runx2, indicative of a mesenchymal (dermal) nature (Abzhanov et al., 2007). Collagen 4 (COL4), elastin, and phosphorylated myosin light chain (pMLC) were detected in both the epidermal and mesenchymal layers. Capillaries beneath the αSMA^+^ layer (and along/on the meninges) were also positive for COL4 and pMLC.

**Fig. 1.**
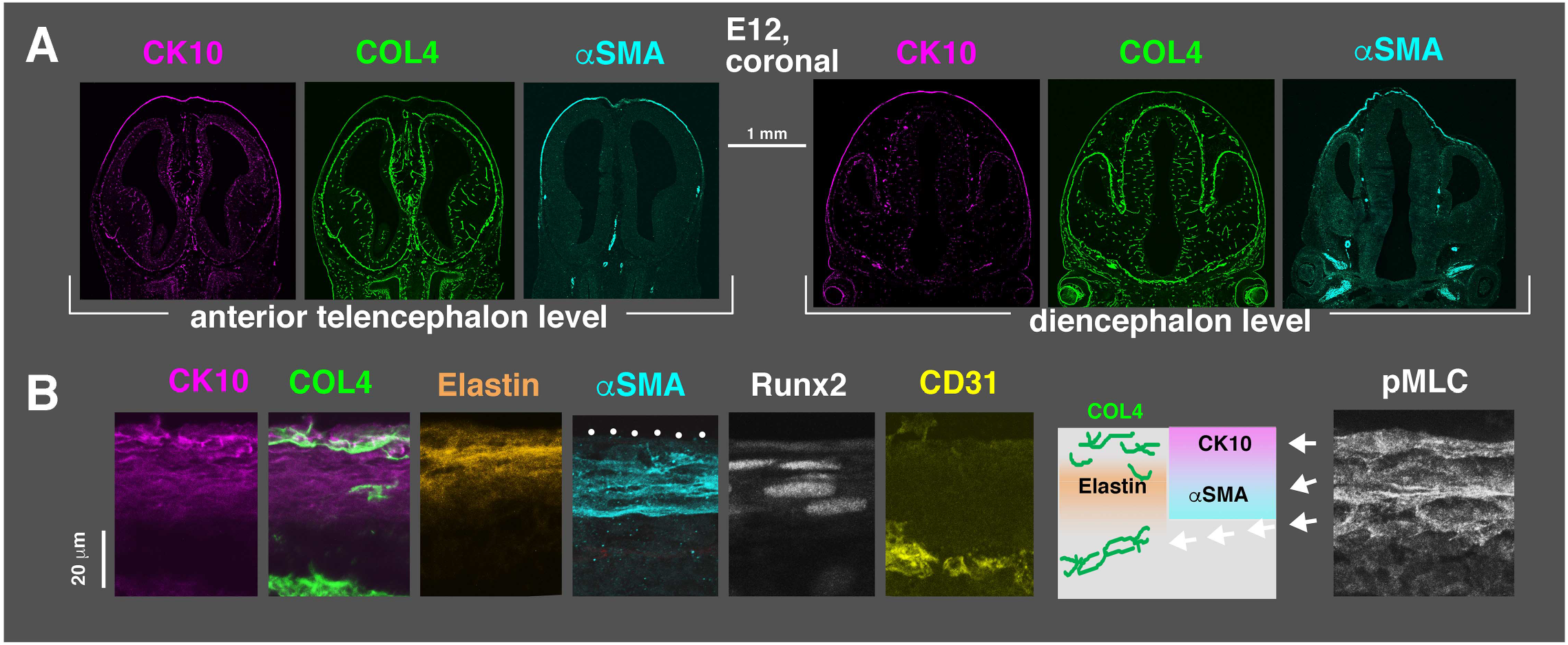
Immunohistochemical characterization of E12 scalp layers. (A) Low-power micrographs of coronal sections immunostained with anti-cytokeratin 10 (CX10), anti-collagen 4 (COL4), and anti-alpha smooth muscle actin (αSMA). (B) Magnified images of the dorsal scalp showing differential or overlapping staining patterns for CX10, COL4, Elastin, and αSMA (schematically illustrated), as well as Runx2, CD31, and phosphorylated myosin light chain (pMLC).

### The E11–E13 scalp was passively stretched and in tension

For execution of a possible constricting or confining action against the brain as hypothesized (Fig. S1B), the scalp/calvarial tissue may have to exert contractility and/or elasticity *in situ*. If such a mechanical confrontation with growing brains occurs, the scalp/calvarial tissue should be in tension. To investigate this possibility, microsurgical tests to release residual tissue stresses were performed on freshly isolated embryos. When a small incision was made to the scalp along the anterior-posterior (AP) axis of the head (both the epidermal and mesenchymal layers were cut, but the underlying meninges were kept intact), the cut edges of the scalp immediately separated along the dorsoventral (DV) axis of the head, and the wound continued to separate for 30 min (n=2/2 at E11; n=5/5 at E12; n=3/3 at E13) (Fig. 2A–C). By contrast, incisions made along the axial (body trunk) structures did not result in wide wound opening along the DV axis (Fig. 2C). A DV incision to the scalp resulted in a similarly wide wound along the AP axis (n=2/2) (data not shown). To determine whether this wound-opening response in the scalp truly reflected a physiological mechanism rather than happening because of embryo isolation and incubation in culture dishes, we made incisions to E13 embryos with beating hearts that were connected via the normal umbilical circulation to their anaesthetized mother mice and found that the wound opening was reproduced (n=3/3) (Fig. 2D). To further quantify the degree of shrinkage (i.e., the recoil from a pre-stretched state) of the E13 scalp, a circular incision was made, and measurements were taken from the top view (Fig. 2E), coronal section view (Fig. 2F), or 3D view (Fig. 2G, Movie 1). Quantitatively, the scalp, when disconnected from the skin of the face, shrunk centripetally to 52±11% (mean±SD, n=5) (top-view analysis) of the original size/area by 20 min after incision and shrunk further to 25±5% (n=3) (3D area measurement by Zephyr) by 40 min. Compared to the Zephyr-based 3D measurement (Fig. 2G, Movie 1), the top-view measurement may have underestimated the shrinkage of the excised scalp because the area associated with a dome-like morphology may have been flat-projected and interpreted to be shorter.

**Fig. 2.**
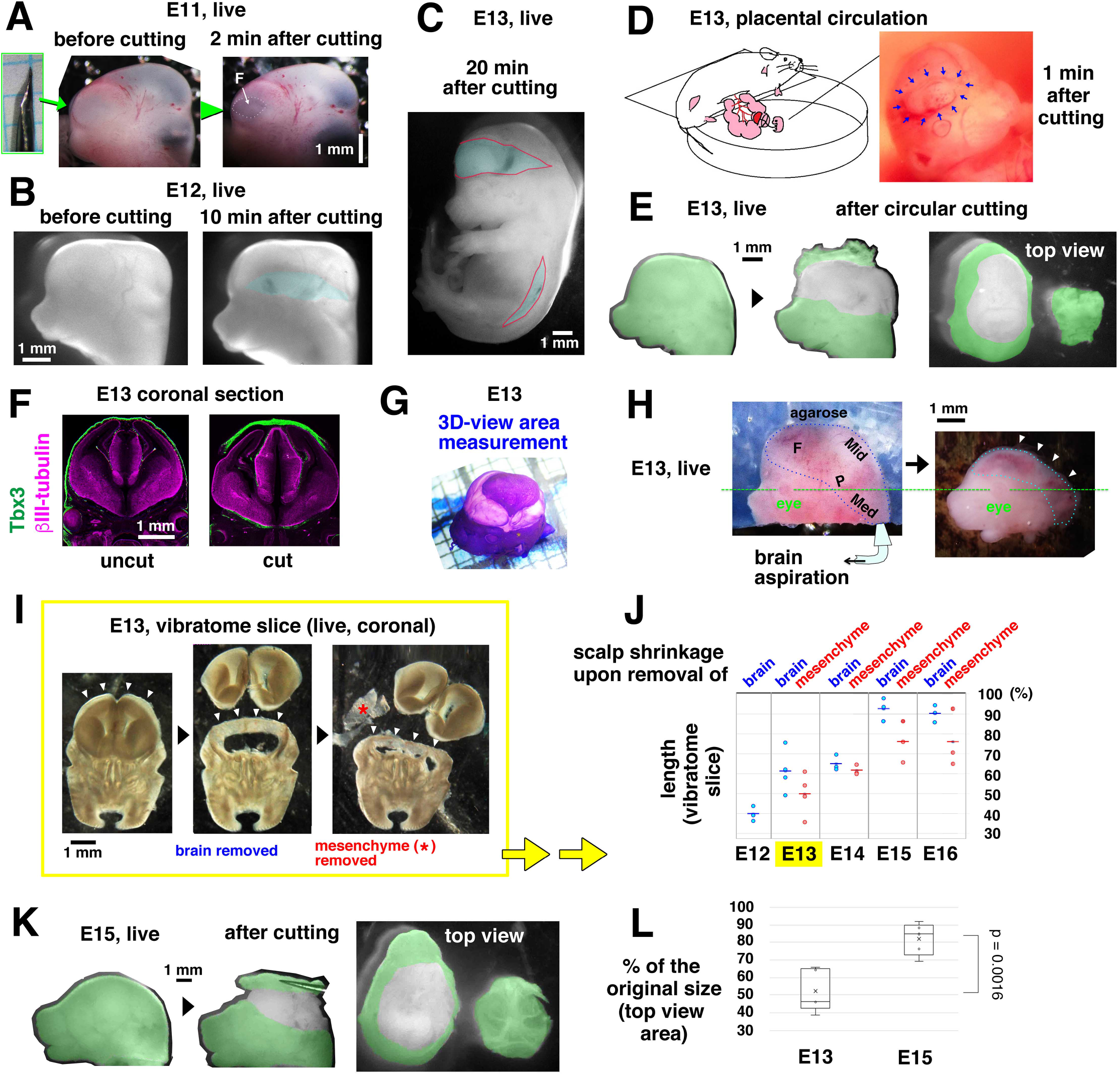
Stress-releasing tests revealed passive stretching and tension in the early embryonic scalp. (A) A horizontal incision to the forebrain (F)-covering scalp of an E11 embryo immediately resulted in a wound. (B) Incision to the scalp of an E12 embryo followed by wound opening. (C) An E13 embryo subjected to an incision in the scalp over the brain and another incision near the spinal cord. (D) Incision made to an E13 embryo with a beating heart that was connected to the anaesthetized mother mouse. The incision produced an immediate wound-opening response. (E) Circular incision and subsequent shrinkage of the scalp at E13. (F) Coronal sections of control and scalp-shrunken heads immunostained for the epidermal marker Tbx3 and the neuronal marker βIII-tubulin. (G) A circularly incised E13 head stained with toluidine blue for 3D area measurement using Zephyr (see also Movie 1). (H) An E13 head freshly mounted in a block made with low-melting-temperature agarose (left), from which the brain (F, forebrain; Mid, midbrain; P, pons; Med, medulla) was then aspirated off (right) until the scalp collapsed (arrowhead). (I) Removal of the brain (cerebral hemisphere) and mesenchyme from a vibratome slice of the head at E13 (see also Movie 2). (J) Graph summarizing stage-dependent changes in the degree of shortening of the scalp upon removal of the brain as well as the mesenchyme (when applicable) from vibratome slices of the head. (K) An E15 head subjected to circular scalp incision. (L) Graph comparing the shrinkage of the circularly excised scalp between E13 and E15.

As another means to test whether the abovementioned brain–scalp mechanical confrontation occurred, we aspirated the brains from E13 heads (Fig. 2H) and found that the dorsal scalp shrunk with a kinetics/magnitude similar to that of the circularly incised scalp (Fig. 2E).

Further analysis of scalp shrinkage upon brain removal using coronal vibratome slices of the head (Fig. 2I, Movie 2) showed a stage dependence that was much weaker at E15~E16 than at E12~E14 (Fig. 2J). When we also removed mesenchymal tissues, which gradually thickened and stiffened between E13 and E15 (probably reflecting differentiation towards osteogenesis [Rice et al., 2003; Roybal et al., 2010]), shrinkage of the remaining scalp (almost purely epidermis) was facilitated overall, especially at E15 and E16 (though it was still weaker than at E12 or E13). A top-view comparison of scalp shrinkage between E15 (Fig. 2K) and E13 (Fig. 2E) showed that the recoil activity of the scalp was significantly greater at E13 (52 ±11% [mean±SD], n=5) than at E15 (82±8%, n=5) (p=0.0016, Welch’s test) (Fig. 2L). These results suggest that the E11–E13 period can be characterized as a stage when the scalp exhibits a remarkable tissue-level responsiveness to recoil when released and that the *in vivo* pre-stretch prerequisite for this recoil depends on the existence of the brain.

### Actomyosin inhibition reduced scalp shrinkage, while calyculin A-mediated contraction of scalp cells caused buckling of the brain wall

To determine whether the pre-stretched *in vivo* situation of the scalp, as evidenced by recoils in response to stress-releasing tests and as found to be mediated by the brain (Fig. 2), can also be explained by autonomous contractility of the scalp given that the scalp was positive for pMLC (Fig. 1B), we pharmacologically inhibited actomyosin (Fig. 3A,B). Shrinkage of the excised E12 scalps was significantly weaker when the scalps were exposed to blebbistatin (a myosin II inhibitor) (shrinkage to 67±6% of the original size [n=7], p=0.018 [Welch’s test]) or Y27632 (rho kinase inhibitor) (to 70±6% [n=7], p=0.007) than in the control (DMSO) group (to 55±12%, n=7).

**Fig. 3.**
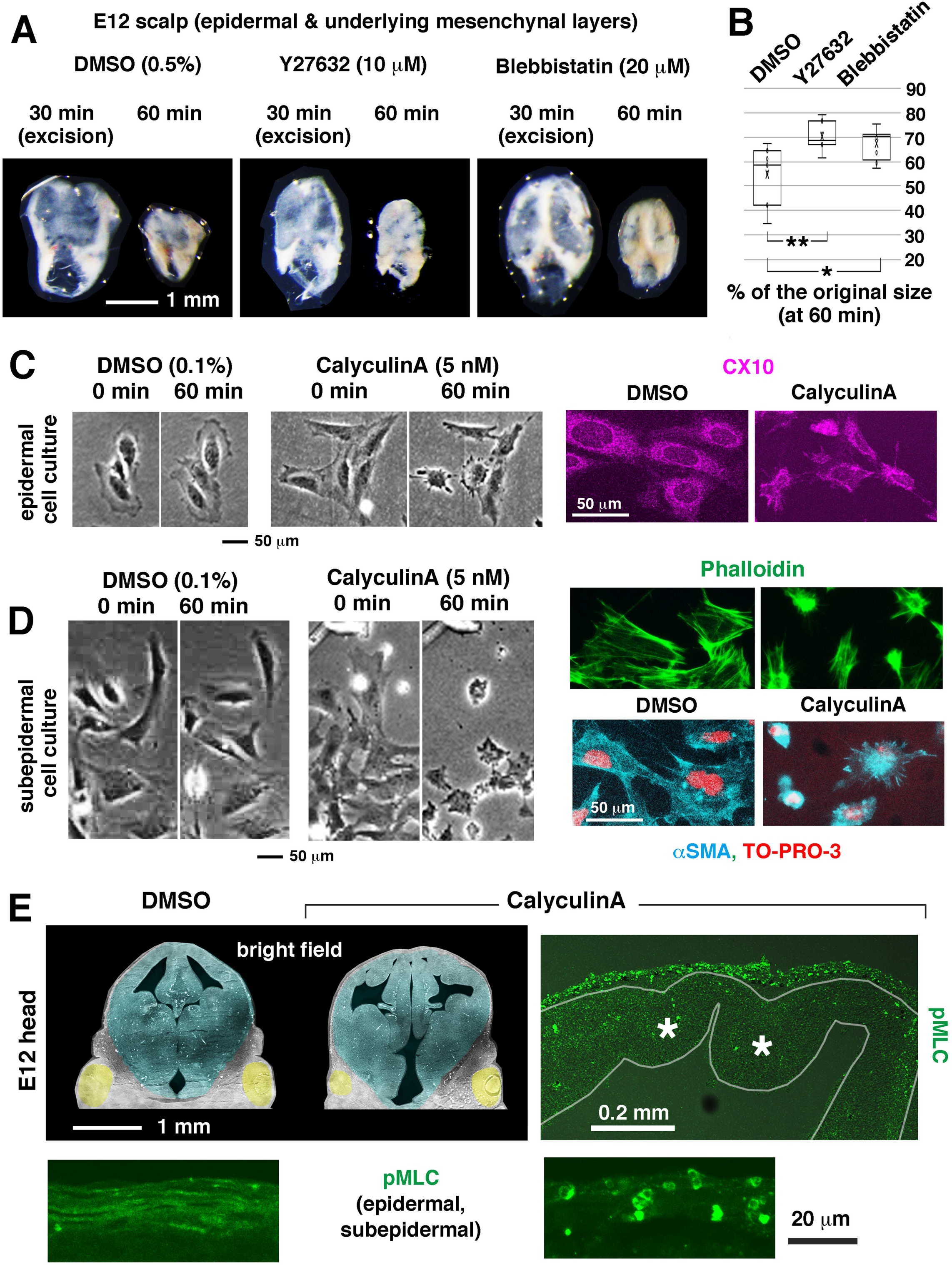
Pharmacological assessment of scalp contractility. (A) Top views of circularly excised scalps under DMSO, Y27632, or blebbistatin treatment. (B) Graph comparing the shrinkage of the circularly excised scalp among the control (DMSO), Y27632, and blebbistatin groups. *, p=0.018. **, p=0.007. (C) Epidermal cells harvested from an E13 scalp and grown for 12 h shrank in response to calyculin A (then confirmed to be of epidermal nature via CX10 immunoreactivity). (D) Cells from the E13 subepidermal layer 12 h in culture also shrank in response to calyculin A (then found to be positive for αSMA). (E) Coronal sections of control (DMSO-treated) and calyculin A-treated E12 heads with high-magnification images of pMLC immunostaining. *, abnormal convexity towards the lateral ventricle.

To next ask whether the hypothetical scalp–brain mechanical confrontation, presumably balanced *in vivo*, is based on optimized myosin-mediated contractility of the scalp, we sought to artificially increase scalp contractility via over-activation of actomyosin. To this end, we took advantage of calyculin A, a phosphatase inhibitor that induces contraction of MLC-or αSMA-expressing cells (Suzuki and Itoh, 1993; Sugita et al., 2019). When separate cultures of epidermal or mesenchymal cells spread flat on the dish surface were exposed to calyculin A, morphological changes (i.e., shortening or rounding) consistent with contraction were observed in each culture (Fig. 3C, D). We then treated E12 heads with calyculin A (1 μM, 60 min) and found in coronal sectional inspections that pMLC immunoreactivity was increased in the scalps with spot-like rather than normal fibrous staining patterns and that cerebral walls were buckled with multiple abnormal convexities towards the ventricle (n=4/4) (Fig. 3E). These results suggest that an optimized level of contractility dependent on actomyosin *in vivo* may be a major force-generating factor that drives tangential narrowing of the scalp.

### The embryonic scalp also responded to elastase but not to collagenase

To examine the possible contribution of extracellular matrix proteins to the tangential narrowing behaviour of the E11–E13 scalp, we also treated excised E12 scalps with elastase or collagenase. Shrinkage was significantly weaker when the scalps were exposed to elastase (shrinkage to 66±9% of the original size [n=10], p=0.03 [Welch’s test]) but not when the scalps were treated with collagenase (to 59±11% [n=9], p=0.270) than under control conditions (to 55±12%, n=7) (Fig. 4A, B). In E12 embryos treated with elastase, dorsal/outward displacement of the midbrain was observed (n=3/3) (Fig. 4C). This result suggests that elastin-mediated elasticity may also play a mechanical role in the embryonic scalp.

**Fig. 4.**
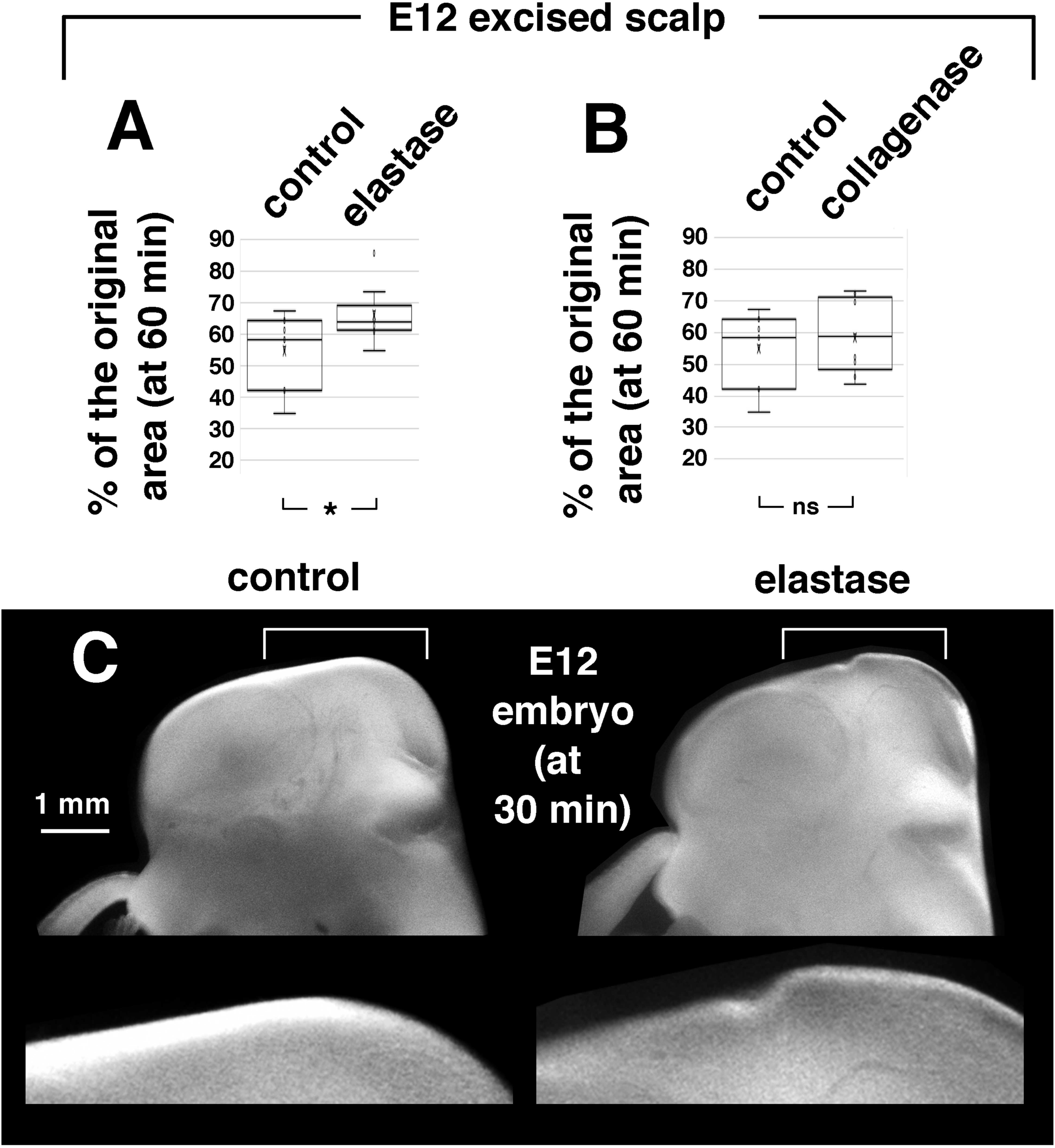
Assessment of the involvement of elastin and collagen in recoil of the excised scalp. (A) Graph showing scalp shrinkage under treatment with elastase (2 U/ml, 60 min in total [30 min after excision]). *, p=0.03. (B) Graph showing scalp shrinkage under treatment with collagenase (2 mg/ml, 60 min in total [30 min after excision]). (C) Left-side views of control and elastase-treated E12 embryos with magnified views of the anterior midbrain region.

### Scalp removal affected brain morphology and cerebrospinal fluid distribution

We next asked whether the dorsal scalp, which is tangentially contractile and elastic, as shown above, truly contributes to confinement of the brain, thereby causing stage-dependent morphological changes. We first focused on the degree of folding of the brain accompanied by narrowing of the fourth ventricle (as shown in Fig. S1B). The angle between the pons and the medulla (P–Med angle) and the angle between the medulla and the cervical spinal cord (Med–S angle) were compared among scalp-intact heads, scalp-removed heads, and isolated brains at E13 (Fig. 5A–C). The P–Med angle was significantly greater in the scalp-removed heads (76±6° [n=7], p=4.31×10^-5^ [Welch’s test]) and the isolated brains (82±6° [n=11], p=1.61×10^-9^) than in the scalp-intact (control) heads (60±6° [n=13]), with widening of the fourth ventricle along the rostrocaudal axis. This observation was consistent with the observation that the brain apparently showed a tendency for dorsal extrusion (similar to the phenomenon seen in an opened jack-in-the-box) in E13 heads whose scalps were circularly incised to induce centripetal recoil (Fig. 2). The Med–S angle was significantly greater in the scalp-removed heads (89±5° [n=7], p=0.0048) but not in the isolated brains (85±5° [n=11], p=0.0516), than in the scalp-intact (control) heads (81±6° [n=10]).

**Fig. 5.**
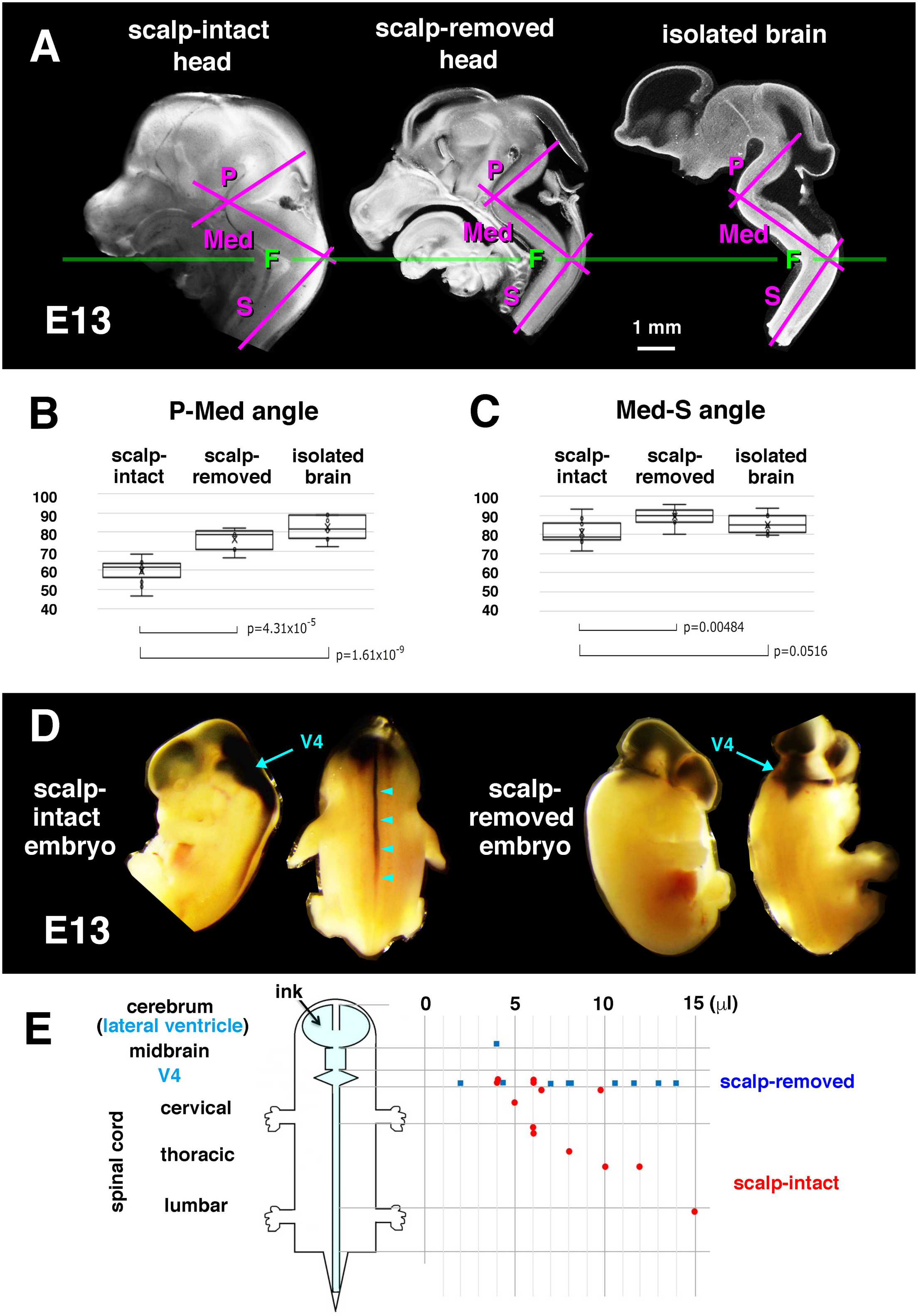
Effects of scalp removal on brain morphology and eCSF homeostasis at E13. (A) Sagittally sectioned right halves of a scalp-intact (normal) head, a scalp-removed head, and an isolated brain. P, pons. Med, medulla. S, cervical spinal cord. F, cervical flexure. (B) Graph comparing the P–Med angle among the three groups. (C) Graph comparing the Med–S angle among the three groups. (D) E13 normal and scalp-removed embryos injected with ink in the lateral ventricle. V4, forth ventricle. Arrowhead, ink spread into the spinal cord. (E) Graph showing the relationship between the volume of injected ink and the extent of ink spread towards the caudal spinal cord in scalp-intact (red) and scalp-removed (blue) E13 embryos.

Since mechanical homeostasis of embryonic cerebrospinal fluid (eCSF) influences, via outward pushing force, the surface morphology of the developing brain (Pexieder and Jelinek, 1970; Desmond and Jacobson, 1977; Garcia et al., 2019), our discovery of this scalp-to-brain confinement prompted us to ask in turn whether the contractile/elastic scalp might assist the brain in its distribution of eCSF. We focused specifically on eCSF delivery to the spinal cord, which has been speculated to occur under the influence of intracranial pressure (Akai et al., 2018). While there are multiple eCSF-producing choroid plexus regions in the brain, the spinal cord lacks them. Inspired by a cardiovascular system example in which the lower leg muscles contract and push the underlying veins to allow them to passively return blood to the heart (Calnan et al., 1970), we hypothesized that the E11–E13 scalp might play a constricting role similar to a leg muscle, contributing to normal eCSF distribution via its compression activity. To test this hypothesis, we injected ink into the lateral ventricle in scalp-intact or scalp-removed embryos (Fig. 5D). In scalp-intact embryos, ink easily spread caudally in a dose-dependent manner, entering the cervical spinal cord at a volume of approximately 5 μl and reaching more caudal levels at 10~15 μl. In contrast, ink injected into scalp-removed embryos stayed in the brain, filling the ventricles in the brain to a great extent without entering the spinal cord (Fig. 5D, E). These results obtained in acute analysis suggest that the scalp may mechanically contribute to morphogenesis and eCSF homeostasis during brain development.

### Scalp cell proliferation was decreased in brain-removed heads, while cells of excised scalps proliferated well when the tissue was re-stretched in culture

Since physiological and clinically oriented skin expansion via cell proliferation is mediated by stretching (Zöllner et al., 2013; Aragona et al., 2020), we next investigated whether passive tangential stretching of the embryonic scalp, as is expected to occur during the mechanical confrontation of the scalp with the outwardly expanding brain *in vivo* (Fig. S1B), is associated with (or contributes to) normal proliferation in the scalp. To this end, we prepared two different 3D culture systems. First, normal and brain-removed heads at E13 were cultured for 4 h, with bromodeoxiuridine (BrdU) application during the final 30 min. Low-power coronal section inspection revealed shrinkage of the scalp in the brain-removed heads (to 50% [on average, n=3] of the brain-intact scalp) (Fig. 6A). Reflecting such tangential narrowing, the CX10^+^ epidermal layer was considerably thickened in high-magnification views (Fig. 6B). The total number of BrdU^+^ nuclei counted in the entire dorsal scalp length (between the bilateral eyelids) was significantly lower in the brain-removed group (54±9 [n=3], p=0.0013 [Welch’s test]) than in the normal group (104±4, n=3) (Fig. 6B, C), suggesting that cells that were in S phase of the cell cycle were significantly more abundant in the stretched epidermis of the brain-intact head than in the shrunken epidermis of the brain-removed head.

**Fig. 6.**
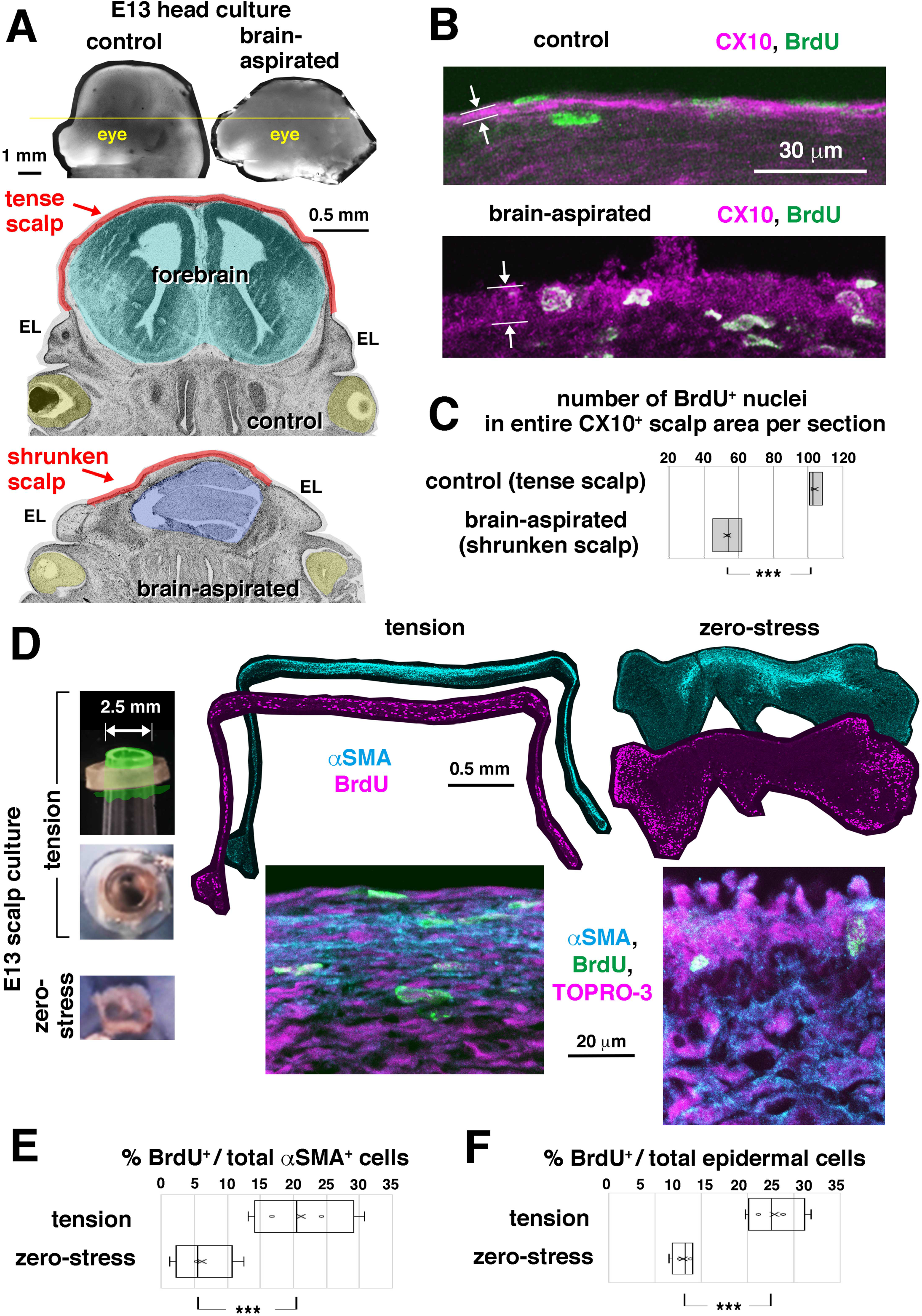
Assessment of the role of brain-mediated mechanical stretching on proliferation of scalp cells. (A) Left-side views and coronal section views (at the level of the eye, highlighted in yellow) of normal and brain-removed E13 embryos cultured for 4 h with bromodeoxyuridine (BrdU) during the final 30 min. EL, eyelid. (B) Magnified photomicrographs of the dorsal scalp immunostained with anti-CX10 and anti-BrdU. The epidermal layer (arrow) was thicker in the tangentially shrunken scalps of the brain-removed heads. (C) Graph comparing the total BrdU^+^ nuclei found in the CX10^+^ epidermal layer of the entire scalp area (defined as the area between the bilateral eyelids). (D) Circularly excised and shrunken E13 scalps were then cultured under re-stretched or zero-stress conditions for 4 h (with BrdU during the last 30 min) before cross sectioning and immunostaining with anti-αSMA and anti-BrdU. A silicone rubber ring was used to tighten each stretched scalp (schematically shown in green in the side-view panel [upper], real picture in the top-view panel [lower]) over the cut surface of a pipette tip. (E) Graph comparing the proportion of total αSMA^+^ cells (visualized with TOPRO-3) that were also positive for BrdU between the two conditions. On average, 125 cells were counted in the tension culture group (n=4) and 96 cells were counted in the zero-stress culture (n=4). (F) Graph comparing the proportion of total epidermal cells (most superficial, αSMA^-^) that were also positive for BrdU between the two conditions. On average, 32 cells were counted in the tension culture group (n=4) and 73 cells were counter in the zero-stress culture (n=4).

Second, we cultured excised scalps under tensile (stretched) or zero-stress (released) conditions. This was necessary for reliable assessment of the effect of stretching on proliferation in the healthy non-epidermal (mesenchymal) layer because oxygen supply to this relatively deep zone compared to the epidermis seemed insufficient in the first (whole-head culture) method. Another advantage of this second approach was that it allowed us to most clearly distinguish physical and chemical factors that could have together arisen from the brain in the first method. After excised E13 scalps automatically shrunk (when released from the pre-stretched state), some were left recoiled, and some were re-stretched using a device, as shown in Fig. 6D (tightened with a silicone rubber ring to form a tympanum on a 2.5-mm-diameter drum). After 4 h of incubation (with BrdU exposure during the final 30 min), low-power inspections revealed healthy αSMA^+^ mesenchymal layers in both groups. In stretch culture, the αSMA^+^ layer was thin, as it was *in vivo*, and the tangential length was up to 4-5 mm, including the 2.5-mm central tympanum area, while the αSMA^+^ layer was only approximately 1.5-2 mm long and much thicker in zero-stress culture than in stretch culture. BrdU labelling (i.e., occurrence of normal entrance into S phase) occurred similarly at the most distal part (originally most ventral, closest to the eyelid) in both explants. Within the central αSMA^+^ layer, however, the BrdU labelling index was significantly lower in the zero-stress group (6.2±4.7% [n=4], p=0.01 [Welch’s test]) than in the stretched group (21.2±7.9% [n=4]). Similarly, the BrdU labelling index in the central epidermal (αSMA^-^) layer was also significantly smaller in the zero-stress group (8.0±1.2% [n=4], p=0.002) than in the stretched group (17.9±3.1% [n=4]) (Fig. 6E, F). Together, these results suggest that tangential stretch of the scalp, as caused by outward pushing by the brain *in vivo*, may promote the proliferation of epidermal and mesenchymal cells at E13.

### Measurement of pressure needed to mechanically re-balloon the shrunken scalp

Finally, we sought to estimate the approximate level of pressure at which (1) outward pushing from the growing brain and (2) confinement of the brain by the scalp may exist in bi-directional balance (Fig. 7A). To measure pressure for this mechanical brain–scalp confrontation, we referred to a previously developed water manometer (Jelinek and Pexieder, 1968) and prepared a system to monitor pressures sufficient to fully inflate the shrunken scalp of an emptied (brain-removed) head of an E13 embryo (Fig. 7B, C). By lifting a medium-containing syringe connected to the embryo’s collapsed head (Fig. 7C, left panel), culture medium was sent to the cranium. By measuring the height to which the syringe had to be lifted (“h” in Fig. 7C, 8~12 mm) to result in apparently sufficient re-ballooning of the scalp to restore the original contour (Fig. 7C, middle panel), we estimated the pressure to be 77±18 Pa (n=19). This value was higher than the eCSF pressure measured at E2~3 in the chick (15~30 Pa) (Jelinek and Pexieder, 1968; Garcia et al., 2019) and lower than the eCSF pressure at E19 in the rat (147 Pa) (Jones and Bucknall, 1987). Since our judgement as to whether the scalp was fully re-stretched to the *in vivo* degree depended only on our monitoring of the contour of the scalp in lateral views, we assessed whether further pressurization might immediately result in excessive ballooning of the scalp. However, we found a lag until clear over-ballooning was detected in the scalp at 93±16 Pa (Fig. 7C, right panel). Therefore, the pressure at which mechanical confrontation between the growing brain and the scalp may occur in E13 mice *in vivo* is estimated to be between 77 Pa and 93 Pa.

**Fig. 7.**
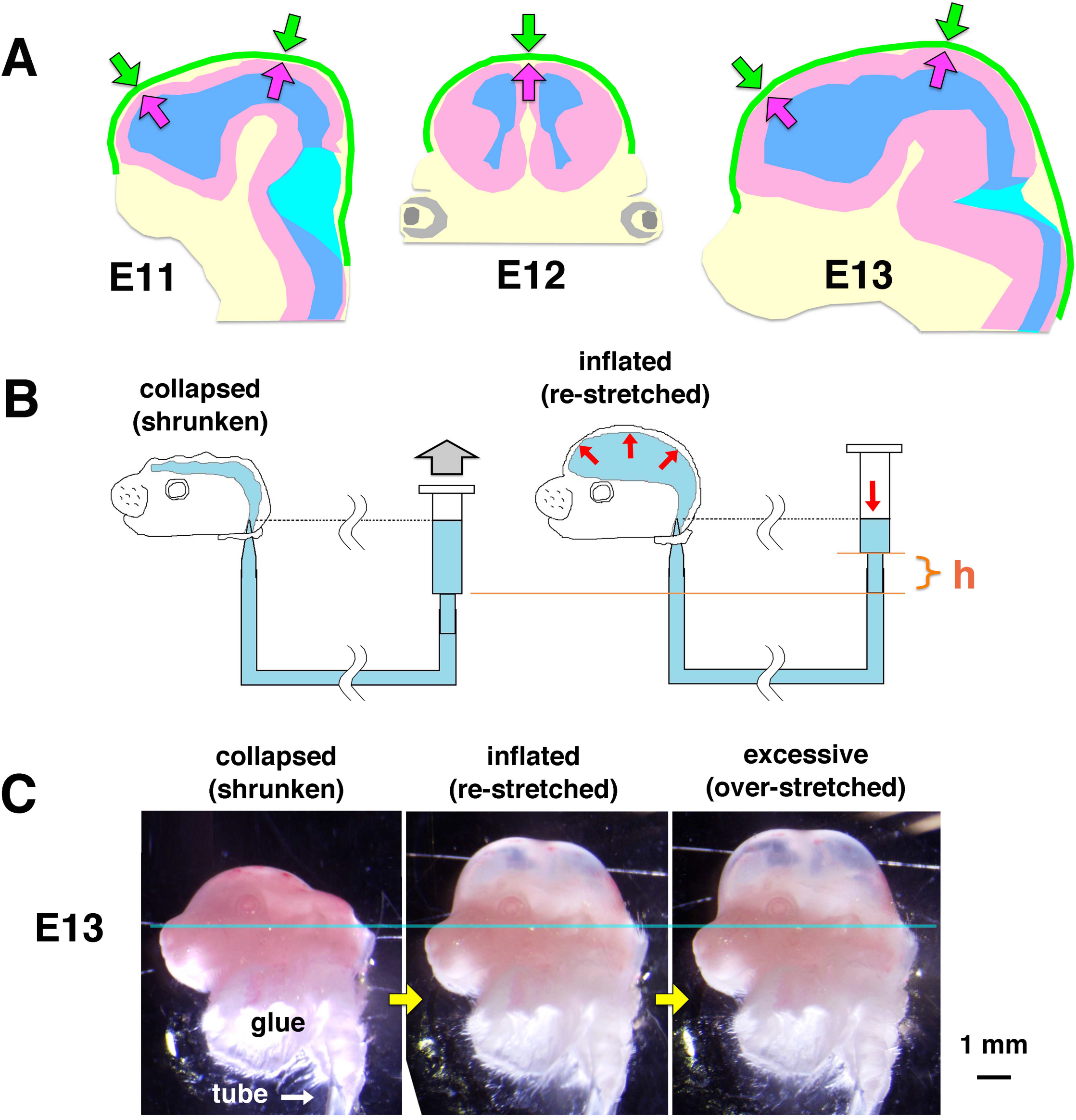
Mechanical confrontation between the brain and the scalp and pressure measurement. (A) Schematic representation of sagittal sections of E11 and E13 heads (from Fig. S1) and a coronal section of an E12 head (from Fig. 6). (B) Schematic illustration of a modified water manometer system applied to the brain-removed head of an E13 embryo for measuring the pressure needed to re-balloon the collapsed/shrunken scalp. A syringe containing culture medium was connected through a flexible tube to the embryo’s collapsed head. The original water surface (in the syringe) and the tip of the tube (in the embryo’s collapsed cranium) were at the same level. When the syringe was lifted to a sufficient height (h), the culture medium sent from the syringe through the tube inflated the cranium to re-balloon the scalp until it regained the original concavity. (C) An E13 head before and after inflation with medium. Blue line, drawn from the lower border of the eye to the ear.

## DISCUSSION

Surgical removal of the scalp has long been no more than a preparative (brain-harvesting) step preceding main research on the developmental mechanisms of the mammalian brain. Surgical procedures such as slicing and incising tissues, in contrast, have been scientifically useful for elucidating mechanical events within the developing brain wall, such as passive cellular movements using a spring-like mechanism based on tissue elasticity (Shinoda et al., 2018), fencing of neural progenitor cells’ moving somata by differentiating cells (Watanabe et al., 2019), expansion of the cerebral area via a landslide-like tissue spreading mechanism (Saito et al., 2019), or abnormal delamination from the apical surface in response to overcrowding (Okamoto et al., 2013). In the present study, however, scalp-targeted surgical treatments revealed remarkable and interesting recoiling of the scalp in E11~13 mice (Fig. 2). Circular incision-mediated centripetal scalp shrinkage continued until the scalp reached 52% of the original size (top view) by 20 min and 25% of its original size (3D view) by 40 min at E13 (Fig. 2). The pre-stretch prerequisite for this cutting-induced recoiling of the scalp was found to depend on (1) optimized levels of actomyosin-mediated contraction in the epidermal and mesenchymal (preosteogenic) layers (Fig. 1,3), (2) elastin-associated extracellular matrix components distributed in these layers (Fig. 1,4), and (3) outward pushing by the growing/expanding brain (at a pressure of approximately 77~93 Pa at E13) (Fig. 2,7). Brain-mediated passive stretching of the scalp may underlie normal scalp cell proliferation (Fig. 6), suggesting brain-to-scalp mechanical assistance. In return, scalp-mediated confinement of the brain may underlie passive brain folding (narrowing of the angle between the pons and the medulla that occurs *in vivo* between E11 and E13) and eCSF homeostasis (Fig. 5), suggesting a scalp-to-brain physical contribution. Thus, the brain and the scalp are in a close relationship to mutually provide mechanical benefits (Fig. 7A). This scalp–brain mechanical collaboration is reminiscent of a recently proposed model of engine-like positive feedback mediated by inter-tissue forces (Xiong et al., 2020; reviewed by Trubuil and Solon 2020), although whether scalp-mediated confinement affects long-term brain-forming events such as proliferation or differentiation remains unknown and needs to be studied.

The buckling of the cerebral wall induced by treatment with calyculin A (Fig. 4) and by hypoxia (Fig. S2), mainly attributable in both cases to over-activation of actomyosin in the scalp and vessel-containing meningeal layers, was morphologically similar to previously observed buckling of cerebral walls during which the Wnt/β-catenin pathway was found to be over-activated to accelerate the proliferation of neural stem cells (Chenn and Walsh, 2002; Okamoto et al., 2013). This suggests a general rule of tissue bucking in which the inner wall/layer can buckle either via overgrowth/proliferation or through external constriction (Nelson, 2016; Trushko et al., 2020). In summary, we combined a variety of microsurgical and measurement methods to reveal a mechanical confrontation and collaboration between the early embryonic mouse brain and the scalp. The techniques used, responses observed at the cellular and tissue levels, and measurement results will be useful for further studies investigating the mechanisms underlying the development of the head, including the brain.

## MATERIALS AND METHODS

### Animals

The animal experiments were conducted according to the Japanese Act on Welfare and Management of Animals, the Guidelines for Proper Conduct of Animal Experiments (published by the Science Council of Japan), and the Fundamental Guidelines for Proper Conduct of Animal Experiments and Related Activities in Academic Research Institutions (published by the Ministry of Education, Culture, Sports, Science and Technology, Japan). All protocols for animal experiments were approved by the Animal Care and Use Committee of Nagoya University (No. 29006). Pregnant female mice (*Mus musculus*) were obtained from SLC (Hamamatsu, Japan; for ICR mice) or by mating at Nagoya University. Embryonic day (E) zero was defined as the day of vaginal plug identification.

### Immunofluorescence

Heads of embryonic mice (E11~E13) and cultured scalps were fixed with periodate-lysine-paraformaldehyde (PLP) fixative (containing 2% paraformaldehyde [PFA]) (4°C 1~3 h). For phospho-myosin light chain (pMLC) immunostaining, heads were fixed in PLP containing 2% trichloroacetic acid (1 h). After immersion in 20% sucrose, the heads were embedded in OCT compound (Miles), frozen and sectioned coronally (16 μm). Cultured cells were fixed in 4% PFA for 15 min (room temperature). The frozen sections or fixed cells were treated with the following primary antibodies: anti-BrdU (rat, NB500-169, Novus Biologicals, 1:2000); anti-βIII-tubulin (mouse, MMS-435P, Covance, 1:1000); anti-CD31/PECAM-1 (goat, AF3628, R&D, 1:300), anti-collagen 4 (rabbit, ab19808, Abcam, 1:500), anti-cytokeratin 10 (mouse, ab9025, Abcam, 1:500), anti-elastin (rabbit, ab21610, Abcam, 1:500), anti–phospho-myosin light chain (pMLC) (rabbit, ab2480, Abcam, 1:500), anti-Runx2 (rabbit, #12556, Cell Signaling, 1:1600), anti-αSMA (rabbit, 23081-1-AP, Proteintech, 1:500), and anti-Tbx3 (rabbit, ab99302, Abcam, 1:300). After washes, the sections were treated with Alexa Fluor 488–, Alexa Fluor 546–, or Alexa Fluor 647–conjugated secondary antibodies (Molecular Probes, A-11029, A-11006, A-11034, A-11030, A-11035, A-11081, A-21236, A-21245, 1:200), and subjected to confocal microscopy (Olympus FV1000, Tokyo, Japan). For cultured cells, Alexa Fluor 546–conjugated phalloidin (A22283, Thermo Fisher Scientific, 1:1000) was used.

### Releasing of residual tissue stress

All surgical procedures for embryos or heads were performed in culture medium (DMEM/F12) under a dissection microscope (Leica MZ7.5 or MZ FLIII). Incisions in the scalps of E11 embryos were made using a micro-knife custom-made from a sewing-machine needle (Fig. 2A). Scalps of E12 and older embryos were incised with micro-scissors (tip diameter 0.05 mm and cutting edge 3 mm, 15000-00, Fine Science Tools). Care was taken not to damage the meninges and brain. For incisions in E13 embryos receiving placental circulation, a pregnant mother was anaesthetized by intraperitoneal administration with a mixture of 0.75 mg/kg medetomidine hydrochloride (Orion Pharma, ZENOAQ), 4 mg/kg midazolam (SANDZ), 5 mg/kg butorphanol tartrate (Meiji Seika), and 1.5 mg/kg atipamezole hydrochloride (Kyoritsu Seiyaku), and myometrium relaxation was induced by intraperitoneal administration of 2 mg/kg ritodrine hydrochloride (FUJIFILM Wako). After midline laparotomy of the mother mouse, the uterine horn was pulled out and dipped into culture medium while the mother was kept on a stage above. Then, a small incision was made in the uterine wall and the amniotic sac to expose the embryo’s head in medium. While surgical procedures were performed on the embryo’s head (scalp), we monitored the heartbeat and placental circulation. Aspiration of brain tissues from the cervical region was performed in culture medium using an aspiration device composed of a gel-loading tip (200 μl, 010-Q, Quality Scientific Plastic) connected through a silicone tube to a suction pump (at 10~50 kPa, SP20, Markos Mefear). Since only a small brain portion could be dislodged and removed by a single aspiration, repetitive aspirations were needed to remove the entire brain.

### Quantitative assessment of scalp shrinkage and brain angles

Heads or excised scalps were imaged using a system composed of a dissection microscope (Leica MZ7.5), an illuminator (SZ2-CLS, Olympus), and a CCD camera (SWIFTCAM, Swift Optical Instruments) or using another system composed of a dissection microscope (Leica MZ FLIII), an illuminator (Hayashi LA150TA), and a CCD camera (ORCA ER, Hamamatsu Photonics). Top-view or cross-sectional analysis of the area of the scalp before or after shrinkage/recoil (Figs. 2-5) was performed using ImageJ (National Institutes of Health, https://imagej.nih.gov/ij/). Measurement of scalp shrinkage through 3D reconstruction (Fig. 2G) was performed after the circularly incised E13 heads were stained with 1% toluidine blue for 10~30 sec (note: pretreatment with 1% Triton X-100 for 5 sec facilitated staining). Approximately 48 images taken while the head was rotated and tilted were reconstructed using 3DF Zephyr (3Dflow, Verona, Italy). The area of the circularly disconnected scalp region (toluidine blue-stained and dorsally shrunken) and the entire area dorsal to the incision line (which represented almost the original scalp area) were compared.

### Pharmacological examinations

After incubating E13 heads for 30 min in culture medium containing 0.5% DMSO (Sigma-Aldrich) only, 0.5% DMSO plus 20 μM blebbistatin (Calbiochem), or 0.5% DMSO plus 20 μM Y27632 (Wako), a circular incision was made on each head. The excised scalps were further incubated under the same conditions for 30 min (total incubation time: 60 min). Cultured epidermal or mesenchymal cells were treated with culture medium containing 0.1% DMSO (Sigma-Aldrich) only or 0.1% DMSO plus 5 nM calyculin A (FUJIFILM Wako). For E12 heads, calyculin A was used at 1 μM for 60 min. Throughout incubation of the heads or scalps, the medium was carefully oxygenized with 95% O_2_/5% CO_2_ to prevent the intra-mesenchymal or peri-meningeal vessels from contracting in response to hypoxia, as reported for embryonic mesenteric regions (Brinks et al., 2016). We indeed observed contraction-like behaviour of the scalp when the heads were incubated under hypoxic conditions, which resulted in buckling of the cerebral walls (Fig. S2).

### Ink injection for evaluating the eCSF system

In each of the scalp-intact or scalp-removed E13 embryos, black ink (SHEAFFER Cartridge 96233, diluted 1:5 in phosphate-buffered saline) was injected into the lateral ventricle of a cerebral hemisphere using a glass capillary needle (created with a Narishige PN-30 tip puller) connected via a silicone rubber tube (1 mm diameter) to a PIPETMAN P200 pipet (GILSON).

### Assessment of scalp cell proliferation

Control and brain-removed E13 heads (Fig. 6A), as well as shrunken and re-stretched scalps (Fig. 6D), were incubated for 4 h in DMEM/F12 containing 5% horse serum (Gibco) and 5% foetal bovine serum (Gibco) in an incubator filled with 45% N_2_, 40% O_2_, and 5% CO_2_ (APM-30D, Astec). Bromodeoxyuridine (BrdU) was added to the culture at 3.5 h, and the heads were fixed in 4% PFA at 4 h. For re-stretch culture (Fig. 6D), each of the excised E13 scalps that was automatically shrinking (released from a pre-stretch *in vivo*) was placed onto the cut surface of a pipette tip (2.5 mm outer diameter) and tightened using a silicone rubber ring (made from a 1-mm-thick silicone rubber sheet [K-125, Togawa Rubber] using hollow punches [63395456, MonotaRO]). Parallel excised scalps that were not re-stretched were cultured under zero-stress (shrunken) conditions.

### Pressure measurement using a water manometer

Modifying a previously developed water manometer system (Jelinek and Pexieder, 1968), we prepared a system to measure pressures sufficient for full inflation of the scalp of an emptied (brain-removed) head of an E13 embryo (Fig. 7B, C). Briefly, a gel-loading tip (200 μl, 010-Q, Quality Scientific Plastic), the distal 25 mm of which was flexible, was inserted into the head with a glue seal at the neck and connected through a silicone rubber tube to a 1 ml syringe (Termo). The syringe, containing DMEM/F12 (specific gravity 1.0168), was set with the tip facing down, and the plunger was removed. The level of the original water surface within the syringe and the level of the tip of the tube in the embryo’s cranium were adjusted to be the same. The embryo’s head was placed in a 60-mm culture dish containing DMEM/F12. Lifting of the syringe to a sufficient height (8~12 mm) sent culture medium into the cranium, expanding the scalp until it regained its original concavity (Fig. 7C, middle) or until it over-ballooned (Fig. 7C, right).

## Acknowledgments

We thank Takeo Matsumoto for valuable instructions about manometers; Namiko Noguchi, and Makoto Masaoka for excellent technical assistance; and members of Miyata laboratory for discussion.

## Competing interests

The authors declare no competing financial interests.

## Author contributions

K. T., K. S. and A.N. carried out experiments, analyzed the data. K.T., K.S. and T.M. designed the study and wrote the manuscript. All authors approved the manuscript.

## Funding

This work was supported by JSPS KAKENHI 16H02457, 19K22683, and 21H02656 (T.M.) and 20K07224 (K.S.).

## SUPPLEMENTARY INFORMATION

**Supplementary Movie 1 (.mp4 file).**
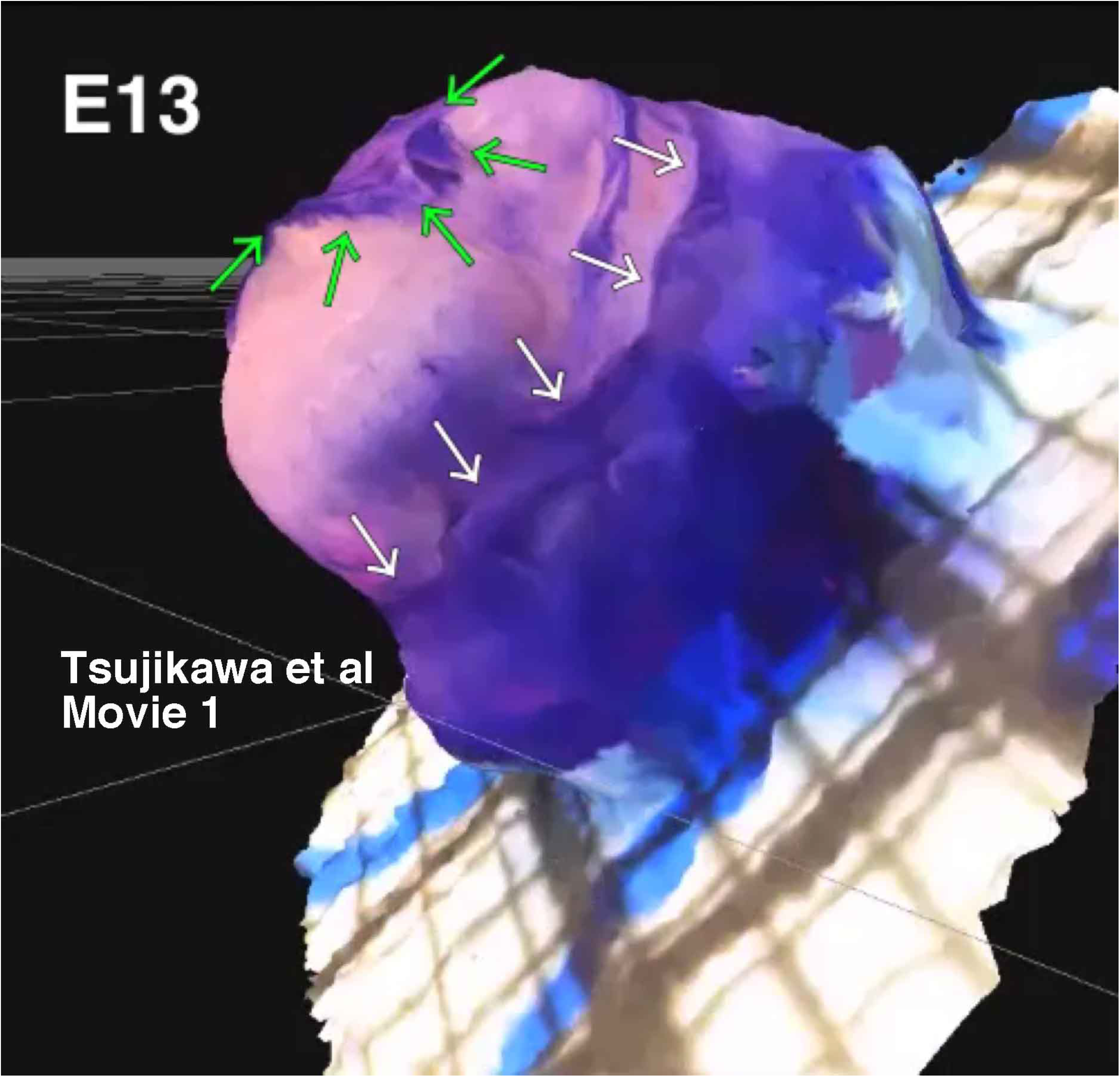
An E12 head whose scalp was circularly incised and shrunk dorsally (green arrows) from the incision line near the eyelids (white arrows). After brief staining with toluidine blue, the head was placed on a stage, and then imaged from different angles for obtaining this 3D reconstruction, which was used for area measurement. Compared to the extensive shrinkage of the distal (dorsal, green arrowed) cut edge, the shrinkage of the proximal/ventral edge was minimal if any.

**Supplementary Movie 2 (.mp4 file).**
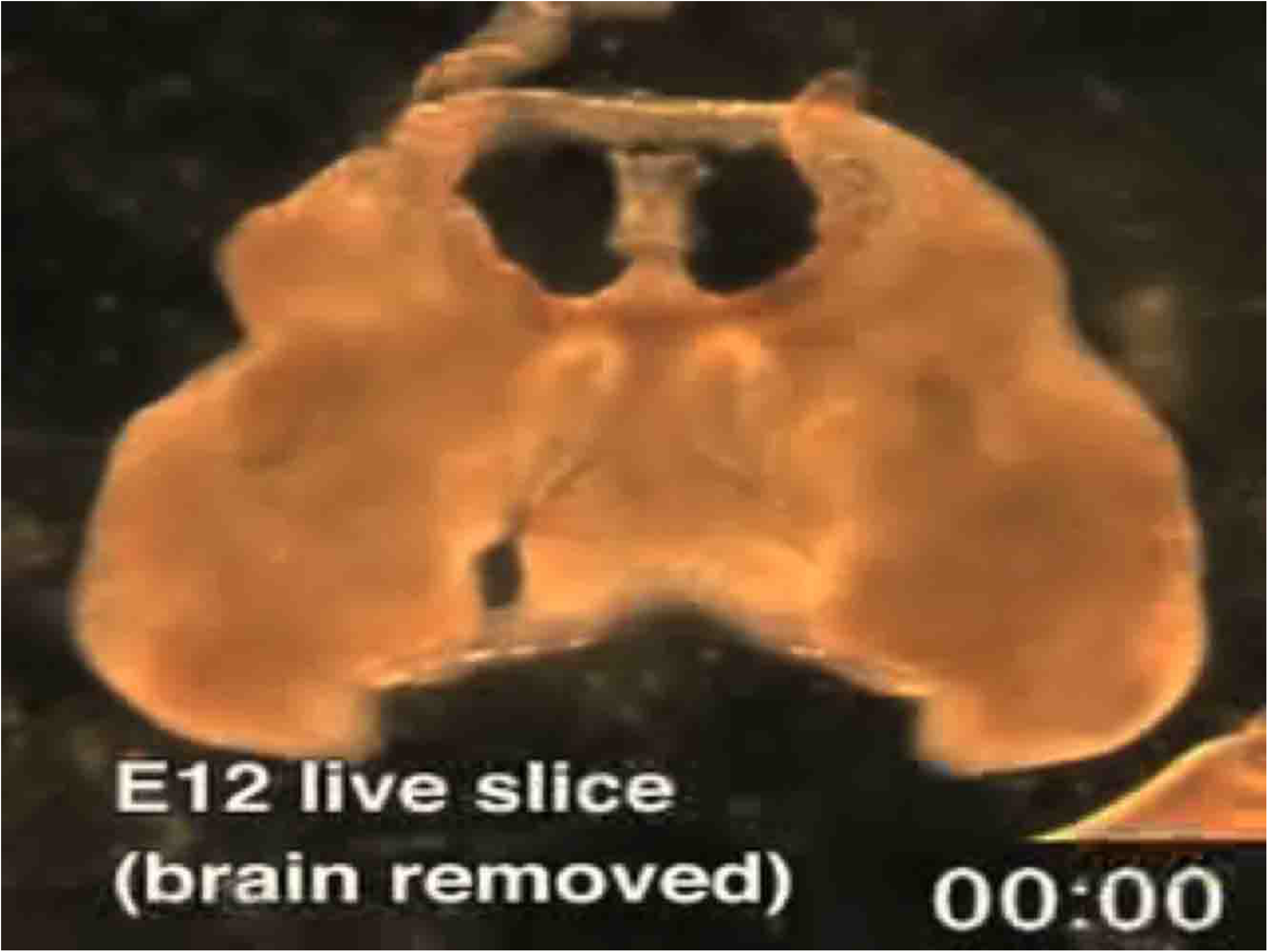
A freshly prepared vibratome slice (0.5 mm) of an E12 head from which the brain had just previously been removed shows shrinkage (i.e., recoil from a pre-stretch) of the scalp (epidermal and mesenchymal layers).

## Supplementary Figures

**Fig. S1.**
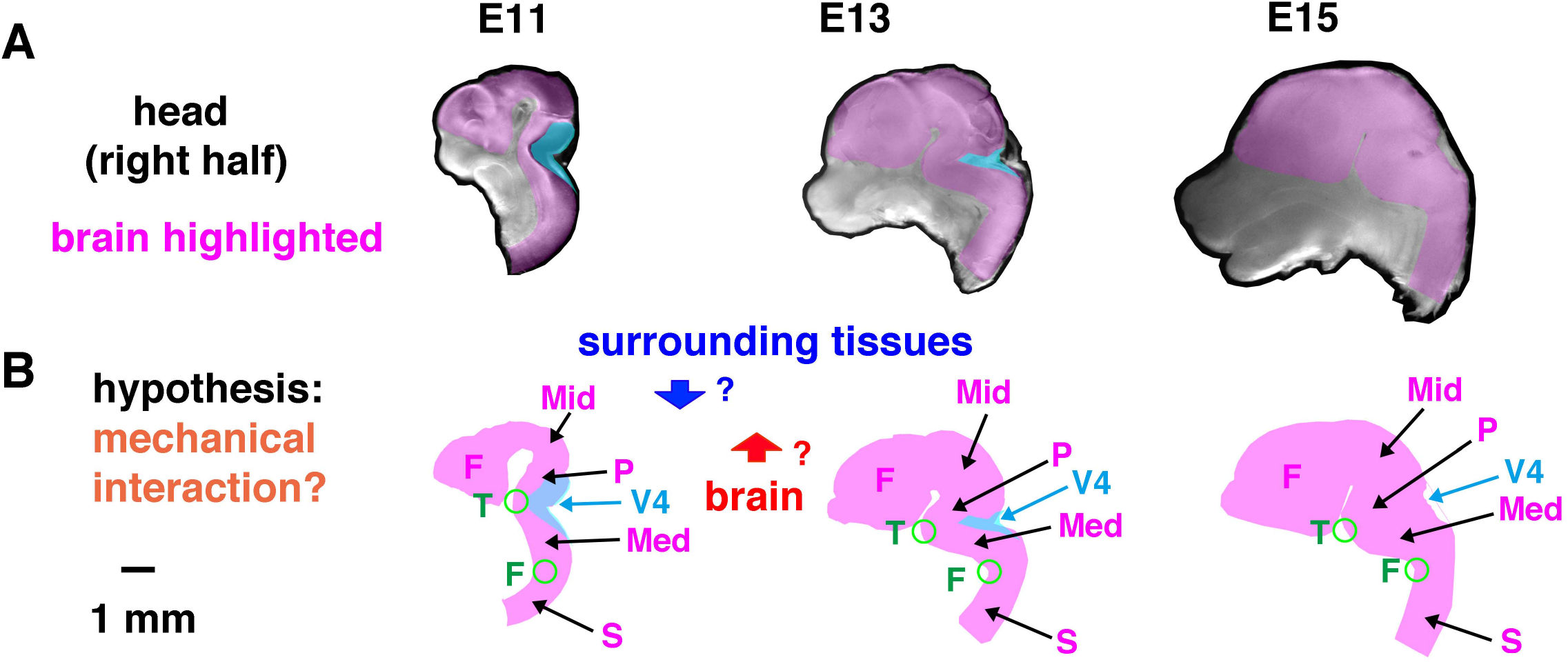
Folding of the pons–medulla axis suggesting possible brain–scalp mechanical interactions during the E11–E13 period. (A) Right half of the head at E11, E13, and E15 (magenta, brain; blue, fourth ventricle). (B) Schematic representations of changing brain morphology. Between E11 and E13, the forebrain (F) and midbrain (Mid) vesicles expanded, the pons (P)–medulla (Med) angle decreased, and the fourth ventricle (V4) narrowed, raising the possibility that the brain pushed and stretched the overlying scalp and that the scalp in return constricted the brain. S, cervical spinal cord. T, trigeminal root. F, cervical flexure.

**Fig. S2.**
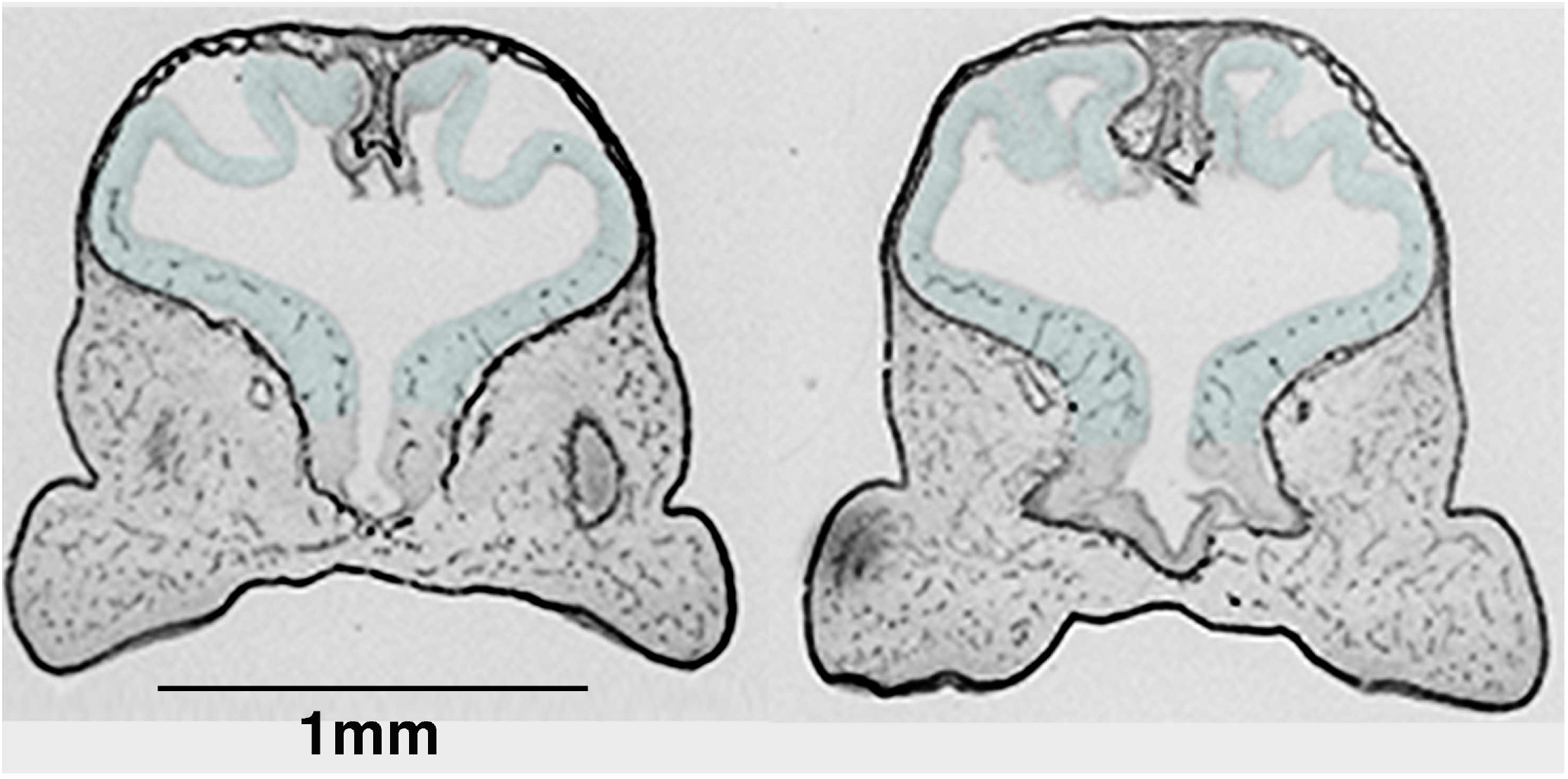
Buckled cerebral wall in an E11 head incubated under hypoxic conditions. Coronal sectional photomicrograph showing an E11 head in which the cerebral wall (highlighted) was buckled. E11 embryos were incubated in a tightly sealed chamber without gas supply or exchange for 60 min. Intra-mesenchymal or peri-meningeal vessels seemed to contract in response to hypoxia via a mechanism reported in capillaries in the embryonic mesenteric region (Brinks et al., 2016).

## Notes

### Competing Interest Statement

The authors have declared no competing interest.

